## Supplementary Material for "3BTRON: A Blood-Brain Barrier Recognition Network"

### Supplementary Information

#### 1 Model Flowchart

Supplementary Figure S1 illustrates the overall workflow of 3BTRON.

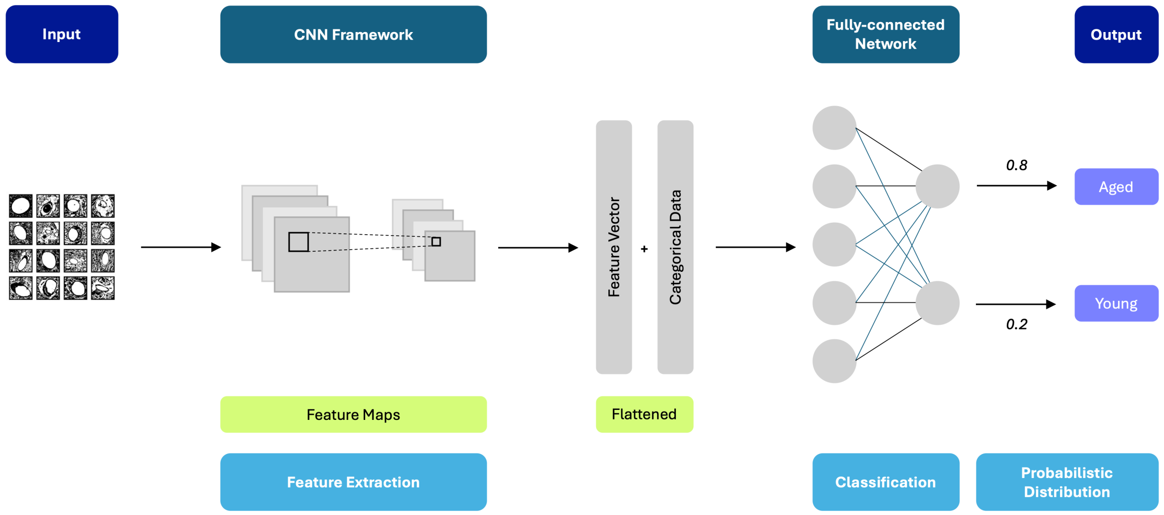


Figure S1: The model framework, illustrating the key steps in the 3BTRON system: image data input, feature extraction using a series of convolutional and pooling layers, generation of image feature vectors combined with sex and brain region data, the fully connected layers, and the output layer/classifier generating binary labels and associated probability estimates.

#### 2 Evaluation Metrics

The blood-brain barrier (BBB) undergoes significant alterations during ageing and the development of age-related diseases, but the full impact on BBB architecture remains unclear. In this work, we have proposed a deep transfer learning framework to identify the BBB architecture, which includes morphological, structural, and textural features, of aged mouse brains. To ensure the both the usefulness and reliability of our model, we assessed the performance of the proposed deep learning models using three evaluation metrics: sensitivity, specificity, and area under the precision-recall curve (AUC Precision-Recall).

Sensitivity (also known as recall) is the proportion of true positive cases correctly identified by the model. It is a measure of how well the model identifies the BBB architecture of aged mouse brains. In contrast, specificity, measures how well the model identifies the BBB architecture of young mouse brains. A high sensitivity indicates that the model is effective at identifying the BBB architecture of aged mouse brains, while a high specificity indicates that the model is effective at identifying the BBB architecture of young mouse brains.

$$Sensitivity = \frac{\mathrm{TP}}{\mathrm{TP} + \mathrm{FN}} \tag{1}$$

$$Specificity = \frac{\mathrm{TN}}{\mathrm{TN} + \mathrm{FP}} \tag{2}$$

where TP, TN, FP, and FN refer to True Positives, True Negatives, False Positives, and False Negatives, respectively.

As aged mouse brains are slightly underrepresented in the dataset compared to young mouse brains, we use the AUC Precision-Recall to optimise the hyperparameters of our model. The AUC Precision-Recall is the area under the curve created by plotting the precision of a model against its recall, measuring how well the model identifies the positive class across all possible thresholds. A high AUC Precision-Recall indicates that the model is effective at correctly identifying the BBB architecture of aged mouse brains while maintaining a low rate of false positives (high precision). Precision is the proportion of correct positive predictions. It is a measure of how confident the model is that the positive prediction is actually correct. A high precision indicates that when the model identifies the BBB architecture of a brain tissue sample as being from an aged mouse, it is likely correct.

$$Specificity = \frac{\mathrm{TP}}{\mathrm{TP} + \mathrm{FP}} \tag{3}$$

The combined use of these metrics helps provide a comprehensive picture of the model's ability to identify BBB architecture of aged mouse brains.

### 3 Evaluation Procedures

As described in the Methodology Section, we chose to evaluate our models using 10-fold cross-validation, before using 5-fold cross-validation to fine-tune decision thresholds (see Figure S2). By using this two-stage cross-validation procedure, we were able to select the best model and optimise for the best possible performance while preventing overfitting.

### 4 Machine Learning Pipeline Evaluated

For this work, we evaluated several combinations of models, as well as the data used as inputs to the models. In this section, we describe the different techniques we experimented with before arriving at our final deep learning pipeline.

We evaluated our network's ability to identify the BBB architecture of aged mouse brains using four pre-trained deep learning models, all based on convolutional network architecture: ResNet50, MobileNetv2, VGG16, and VGG19.

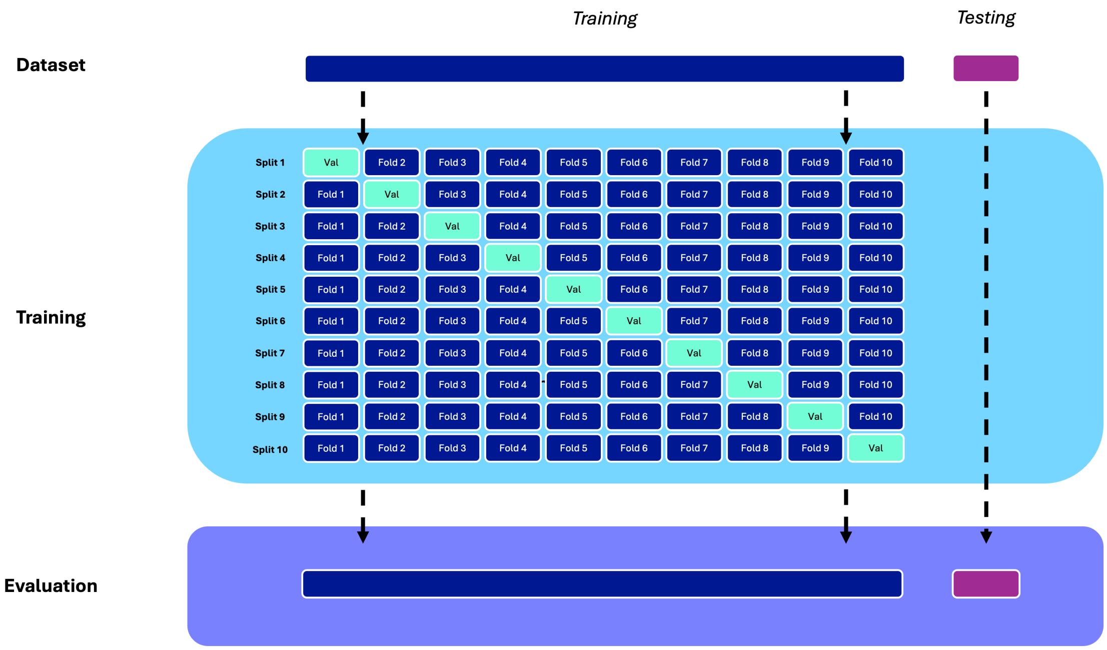


**(a)** Hyperparameter Tuning

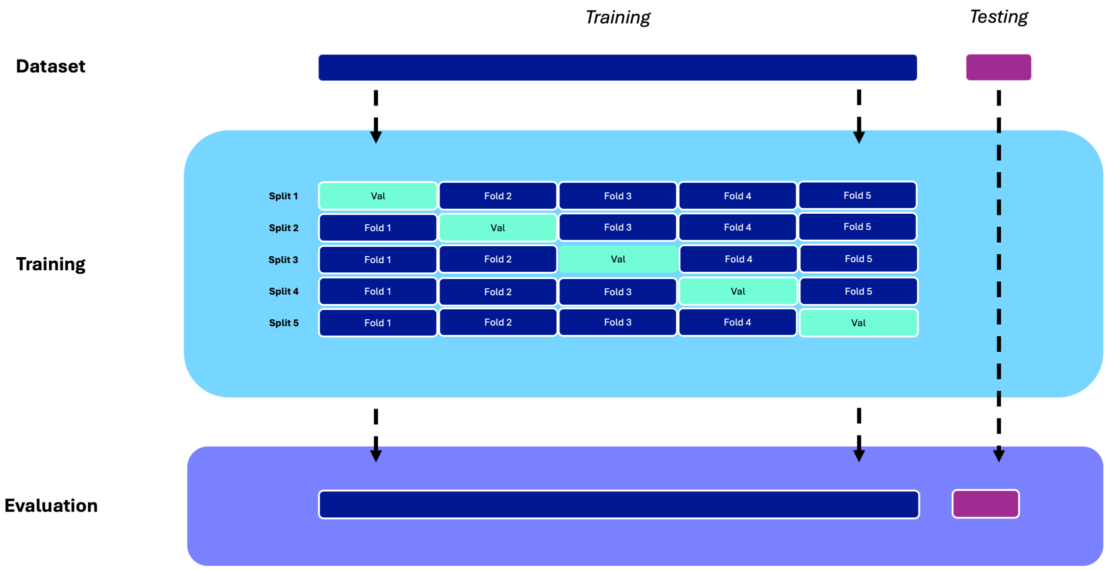


**(b)** Optimising thresholds

Figure S2: The training and validation procedures visualised. a) “Hyperparameter tuning” using 10-fold cross-validation to train the model and evaluating on a held-out test set. b) “Optimising thresholds” using 5-fold cross-validation to train the model and evaluating on the held-out test set.

For all models, we used optimisation to determine the best values of the following hyperparameters:

- batch size, with a value defined in [16, 32, 64, 128, 256]
- maximum number of epochs, with a value defined in [5, 10, 20]
- optimiser type of Adam or SGD
- momentum (only relevant for an SGD optimiser) with a value defined in [0.9, 0.925, 0.95, 0.975]
- learning rate, with a value defined in [1e-7, 1e-2]
- L2 regularisation, with a value defined in [1e-7, 1e-2]

These models were run using Python and implemented using PyTorch.[40] The final model was implemented using the the average number of epochs used for training across the 10-fold cross-validation of the best-performing hyperparameters.

### 5 Reliability and Calibration

In this section, we present the results of the final model's reliability and calibration analysis. Supplementary Figure S3 shows a multi-faceted summary of model performance. The plots on the left-hand side of the figure are reliability plots, showing how well the predicted probabilities of the model align with the true labels (aged and young). In the top plot, we observe some gapping between the average accuracy and the confidence for a given bin, as well as a distance between the average accuracy and average confidence in the lower plot. When considering only positive predictions as in the bottom right-hand plot showing how well the predicted probabilities of the best correspond to the true likelihood of the positive case, we observe that the model was at times overconfident. Finally, the plot on the top right-hand side of the figure shows the distribution of the predicted probabilities made by the model on each class, and here we observe that the model was uncertain about a number of images from young mouse brains, often falsely predicted them as aged based on the application of binary thresholds. These results were a key motivation for the use of stratification, to ensure that the number of false positives could be controlled.

### 6 Stratification Analysis

Previous work by the group reported a stratification analysis of UTI risk.[43] Here, when reporting our predictions, we apply this same approach, stratifying our results into three groups: Green, Amber, and Red, where each group represents low, medium, and high likelihood of age, respectively.

For each split of a 5-fold cross-validation, after training the best performing model on the training data, age likelihood scores are calculated on the validation data so that they can

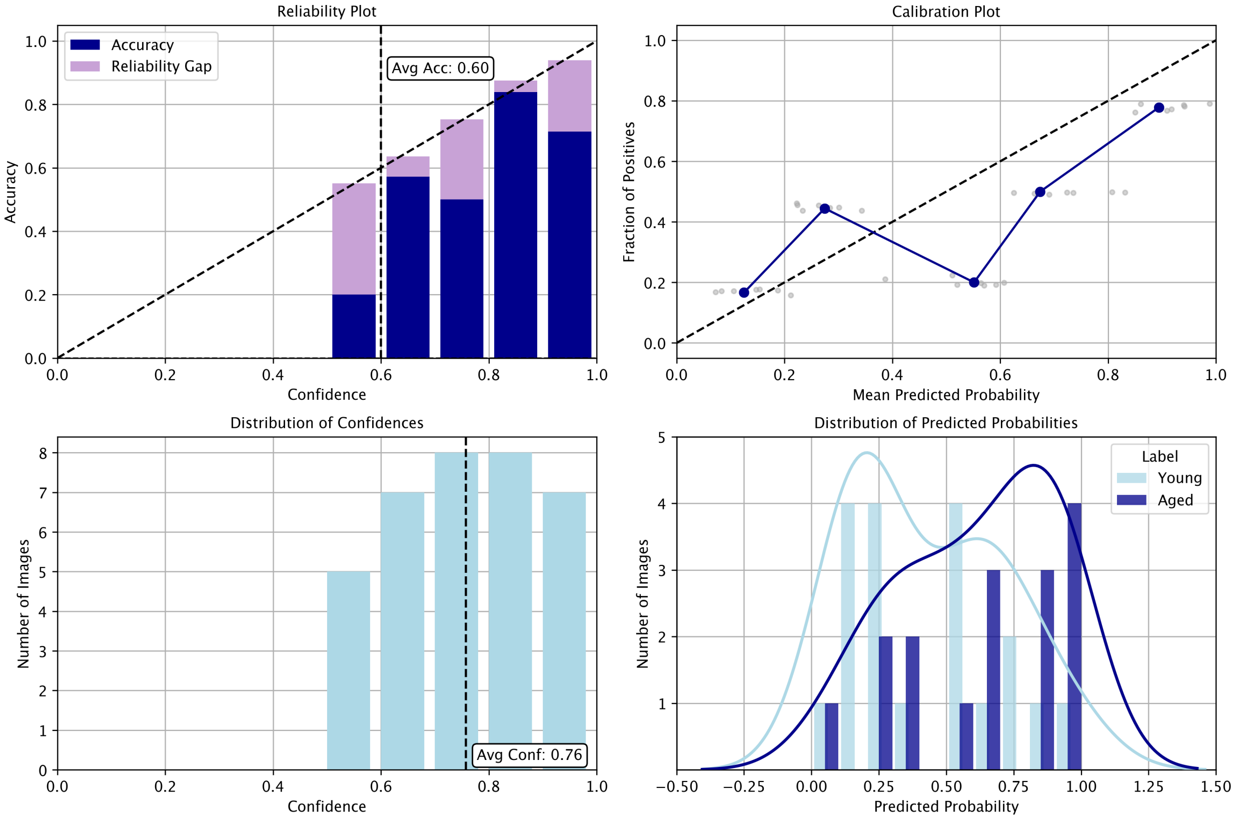


Figure S3: Multi-faceted Summary of Model Performance. The top left-hand plot shows the model confidence (on all images) against accuracy on the held-out test set, with the gaps showing the difference between average accuracy and confidence for a given bin, which would ideally be 0. The bottom left-hand plot shows the histogram of confidences reported on the held-out test set. The top right-hand plot shows shows the mean predicted age likelihood (grouped into 5 bins) against the proportion of aged images for the best model plotted for the held-out test set. Individual data points are depicted in dark grey. The bottom right-hand plots the distribution of probabilities made by the model for each age group.

be used to determine the best thresholds for the final model. To improve the probabilistic output of the model and to reduce any potential overconfidence the model might exhibit, we then applied sigmoid calibration to the predicted probabilities. We then calculate sensitivity, specificity, and precision under different threshold values.

When investigating thresholds, our goal was to maximise sensitivity and specificity while trying to keep model uncertainty (i.e. the number of samples for which the model is not confident about its predictions) low. To achieve this, we varied thresholds with a resolution of 5%, computing these evaluation metrics on the Green and Red groups (negative and positive class groups, respectively) for each pair of thresholds. These thresholds were optimised using the following criteria:

- Ensure that the maximum difference between the green and amber thresholds was no more than 50%.
- Ensure that the range of the green and red groups is equal.
- After employing the previous two restrictions, to jointly maximise sensitivity and specificity by optimising for the J Youden's statistic:

$$J = (Sensitivity + Specificity) - 1 \quad (4)$$

- Finally, we find the thresholds that minimise the difference between sensitivity and specificity.

After employing these restrictions across each split, we found that the best thresholds to apply to the age likelihood score (with range [0, 100]) were as follows: Green $\epsilon$ [0, 25], Amber $\varepsilon$ [25, 75], and Red $\varepsilon$ [75, 100].

#### 7 Performance on Female/Male Split and Different Brain Regions

In this section, we present the model performance on images from female and male mouse brains and across the three different brain regions (corpus callosum - CC, hippocampus - HC, and prefrontal cortex - PFC). We do this using a binary threshold (where the positive class has a threshold $>$ 50% and the negative class has a threshold of $<=$ 50%) and our chosen thresholds (Green $\epsilon$ [0, 25], Amber $\varepsilon$ [25, 75], and Red $\varepsilon$ [75, 100]) to split the predictions made by the proposed model over each demographic and brain region on the held-out test set, calculating the accuracy before and after stratification, as well as the likelihood of a positive prediction. We also report the total number of images for each sex and brain region, as well as the ratio of young and aged images in each demographic (see Supplementary Tables S1 and S2).

Applying binary thresholds highlighted a discrepancy in the accuracy according to both the sex of the mouse from which the image came, as well as the brain region. The discrepancy is particularly noticeable across brain regions, due to the difference in number of aged and

Table S1: Accuracy on images from male and female mouse brains on the held-out test set. We report this before and after stratification is performed. We also show the ratio of young and aged images in each demographic and the likelihood of a positive prediction.

**Before Stratification**

| Sex | Accuracy | No. of Images | Aged : Young | P(Y = 1 \| Sex) |
|---|---|---|---|---|
| **Female** | 64.7 | 17 | 1 : 1.125 | 50.9 |
| **Male** | 55.6 | 18 | 1 : 1.25 | 52.3 |

**After Stratification**

| Sex | Accuracy | No. of Images | Aged : Young | P(Y = 1 \| Sex) |
|---|---|---|---|---|
| **Female** | 77.8 | 9 | 1 : 1.25 | 48.9 |
| **Male** | 80 | 10 | 1 : 1 | 52.8 |

young images for each region. After stratification, we see that accuracy has increased for each sex and brain region, with reduced discrepancy observed between factors (particularly brain regions).

When using this as an investigative tool, researchers should consider how the proportion of labels might impact the metrics. Applied in different research scenarios, providing there is sufficient data, a separate model could be trained to make predictions for each demographic and brain region. However, for this proof-of-concept study, training a separate model for male and female mice and different brain regions would have led to overfitting in the model due to insufficient data.

This experiment also highlighted the advantage of tuning age likelihood score thresholds on individual sexes and different brain regions based on a chosen fairness metric for a specific research scenario or setting. For example, Supplementary Tables S1 and S2 show the likelihood of a positive prediction ($Y = 1$) by sex ($P(Y = 1 \mid Sex)$) and brain region($P(Y = 1 \mid BR)$) using our proposed model with binary thresholds and chosen thresholds. This is referred to as demographic parity, and we see here that our model has high demographic parity across mice of different sexes before and after stratification as the likelihood of a positive prediction is close to equal. While stratification does not fully improve statistical imparity, it does reduce the gap in likelihood between the corpus callosum and hippocampus, as well as significantly increasing accuracies for both the corpus callosum and hippocampus, making the stratification beneficial overall, although the trade-off in fairness should be considered and ideally both accuracy and fairness optimised.

Table S2: Accuracy on images from corpus callosum (CC), hippocampus (HC) and prefrontal cortex (PFC) on the held-out test set. We report this before and after stratification is performed. We also show the ratio of young and aged images in each brain region and the likelihood of a positive prediction.

**Before Stratification**

| Brain Region | Accuracy | No. of Images | Aged : Young | P(Y = 1 \| BR) |
|---|---|---|---|---|
| **CC** | 47.1 | 17 | 1 : 1.43 | 64.3 |
| **HC** | 57.1 | 7 | 1 : 1.33 | 41.3 |
| **PFC** | 81.8 | 11 | 1 : 0.83 | 38.5 |

**After Stratification**

| Brain Region | Accuracy | No. of Images | Aged : Young | P(Y = 1 \| BR) |
|---|---|---|---|---|
| **CC** | 75.0 | 8 | 1 : 0.60 | 66.4 |
| **HC** | 75.0 | 4 | 1 : 3.00 | 50.1 |
| **PFC** | 85.7 | 7 | 1 : 1.33 | 33.9 |
